# Microbial Fuel Cell coupled Microbial Electrolysis Cell for Biohydrogen Production

**DOI:** 10.1101/2025.07.30.667648

**Authors:** Ransford Kingsley Asiedu, Rejoice Ayeley Ayi, Gilbert Blah Quarshie, Michael Kweku Edem Donkor

**Author notes:** **Corresponding Author:** Ransford Kingsley Asiedu.

## Abstract

Hydrogen, a source of renewable energy, unfortunately relies largely on fossil fuel technologies for its production. However, recent studies have shown that microbial technologies could be used to facilitate green hydrogen production. Based on these findings, this work primarily focused on utilizing farm soil, wastewater, anaerobic sludge and cow dung for the production of green hydrogen. A double chambered Microbial Electrolysis Cell (MEC) was coupled with a single chambered Microbial Fuel Cell (MFC) for this work. In this experiment, the fuel cell produced an average potential difference of 118.9 ± 0.001 mV over a period of 312 hours (13 days). The electrolysis cell also produced an average potential difference of 56.8 ± 0.003 mV over the same period of time. The two chambers of the MEC which facilitated the electrolysis process were separated by a locally made Proton Exchange Membrane. Hydrogen gas was produced at an average rate of about 6.9 ± 0.012 mL/day with the highest being 65.5 ± 0.012 *mL* on day 9. Also, an average current of 0.22 ± 0.006 μA flowed through the entire system and the total amount of hydrogen gas produced at the end of the experiment was approximately 96.8 ± 0.012 *mL*. Based on this work, it is evident that green hydrogen can be produced by means of coupling microbial electrolysis cells with microbial fuel cells and utilizing farm soil, wastewater, anaerobic sludge and cow dung.

## Introduction

Reduction in the use of fossil fuel as an energy source is a great way to resolve the alarming issues of global warming and the all-time increasing emissions of carbon. To do this, there is the need to exploit more of environmentally friendly and renewable energy sources (Abdelsalam et al., 2023). Among the several renewable sources of energy, hydrogen stands out as one of the best and efficient sources of energy. Hydrogen offers great potential as an ecofriendly source of energy. It has a wide range of uses and applications. For instance, it is used for the synthesis of ammonia and nitrogenous fertilizers, the production of Rocket fuel and propellant, as a power source for vehicles, the production of fuel cells, Petroleum Refining among many others. However, the means through which we obtain hydrogen is not sustainable, and is environmentally degrading. According to Gencer (2021), about 95% of the world’s hydrogen is generated by fossil fuel-based technologies. This in turn produces about 830 million tons of CO_2_ each year in exchange for 74 million tons of hydrogen. The vast majority of the hydrogen is produced through partial oxidation of heavy hydrocarbons and the steam reforming of natural gas and other light hydrocarbons, otherwise known as Natural Gas Steam Reformation (Rapier, 2020). This is a process within which steam and natural gas are caused to react together under high temperatures and pressures in the presence of a nickel catalyst to produce Hydrogen (H_2_) and Carbon dioxide (CO_2_). This process is not a sustainable means of producing hydrogen as it depends on fossil fuel which is an exhaustible and harmful energy resource. The process also produces lots of waste heat which could be used as a source of energy for other applications. Aside Natural Gas Steam Reformation, there are other popular means of hydrogen production such as Water Electrolysis. This is a process whereby electricity is passed through water to break it down into hydrogen and oxygen. Although the by products are not harmful to the environment, the means through which the electricity is produced raises a huge concern. Water electrolysis is a very expensive way of producing hydrogen as at least 39.4 kWh of electricity is required to produce 1 kg of hydrogen at full conversion efficiency (Tashie-Lewis & Nnabuife, 2021). There are many other fossil fuel-based technologies used to produce hydrogen such as, Coal Gasification, and Plasma Pyrolysis. All these methods leave a huge carbon footprint on the environment. As a way to alleviate the impact of carbon, Microbial Fuel Cells (MFCs) and Microbial Electrolysis Cells (MECs) have emerged as a promising means for the production of biohydrogen. An MFC is an electrochemical cell which generates electricity by utilizing electrons obtained from a biochemical reaction which is propelled by bacteria whereas an MEC is a cell which uses electrochemically active bacteria to transform organic matter into hydrogen and other by-products without contaminating the environment. When combined, these systems create a sustainable way of producing Hydrogen. This is beneficial because both mechanisms are organic and can be easily obtained from waste water treatment plants, garbage facilities, and even refuse dumps. The only limitation is that bacteria work on organic matter and not inorganic matter, so only organic waste can be converted into hydrogen. The MFC produces electrical energy for the MEC which is then used to carry out electrolysis in waste water. This process presents a sustainable way of producing hydrogen that depends on a renewable energy source. The goal is to produce hydrogen gas in a sustainable way, significantly reducing the carbon footprint left behind by fossil fuel-based methods of producing hydrogen and generally to reduce the impact of global warming through innovative technologies.

## Materials and Methods

### Materials and tools

Two 1.5 L and one 5.3 L plastic containers, magnetic stirrer and heater, fibre glass mesh, balloons, epoxy glue, male T connector, salt, aluminium foil, cow dung, oven, citric acid powder, polyvinyl alcohol (PVA) powder, wastewater, activated carbon sponge, carbon felt, farm soil and two multimeters.

### Preparation of PEM

According to the set chemical ratios in do Nascimento et. al. (2020), 2 g of PVA was dissolved in 80 mL of distilled water in a beaker. 1 g of citric acid was then dissolved in the solution and the mixture was stirred for 2 hours to completely dissolve the citric acid using the magnetic stirrer and maintaining a temperature of 80°C. After the citric acid was completely dissolved in the solution, a resin was formed. The resin was then spread on both sides of the fibre glass mesh and the product was wrapped with aluminium foil. It was baked in an oven for 45 minutes at a temperature of 130°C. The resulting PEM is shown in Figure 1.

**Figure 1.**
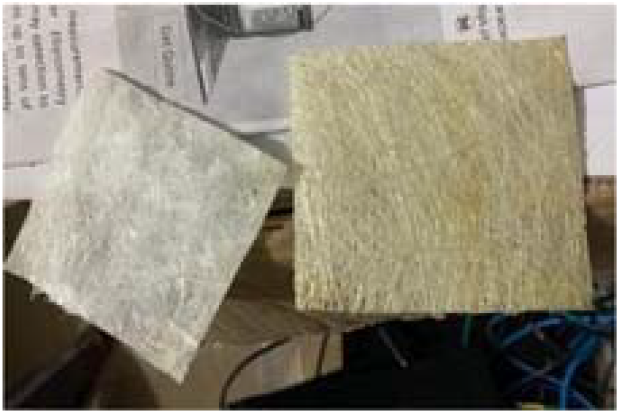
(An image of the locally made proton exchange membrane)

### Construction of the MFC

Farm soil, anaerobic sludge and cow dung were mixed with waste water. The mixture was poured into the 5.3 L plastic container to a height of about 5 cm and the anode (, 22 ×16) was placed on it. The mixture was poured onto the anode to a height of about 10 cm and the cathode (, 22 × 16) was placed on it. The cathode was exposed to oxygen by creating a hole on the lid of the container. Copper wires were passed through the electrodes to allow electron transfer. Figure 2 illustrates the completed single chambered MFC.

**Figure 2.**
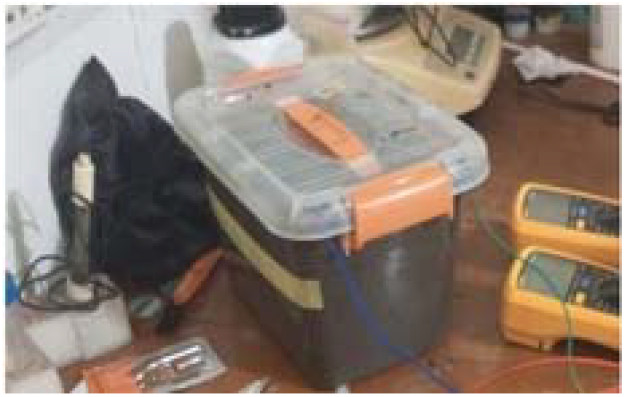
(An image of the constructed single chamber microbial fuel cell)

### Construction of the MEC

A hole was created at one side of each of the two 1.5 L plastic containers. The male T connector was fit into the holes that were created, with the PEM tightly fit to one of its ends. The thick mixture of farm soil, cow dung, anaerobic sludge and wastewater was poured into the anode chamber to a height of about 5 cm and the anode (, 10 × 10) was placed on it. Little of the remaining mixture was poured onto the anode and more of the wastewater was poured onto it to the brim, after which this chamber was covered. The other chamber was filled to the brim with salt solution and the cathode (, 2 × 9) was placed in it. The cathode chamber was then sealed and ensured to be air-tight with only one escape, which was the passage in a hard tube, fixed to the top of the cathode chamber. A balloon was attached to the other end of the hard tube for the collection of the gas. Figure 3 shows the two chambered MEC with the PEM separating it.

**Figure 3.**
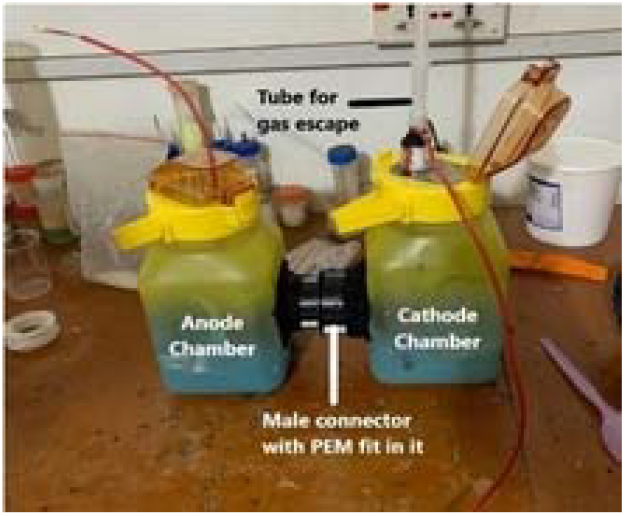
(An image of the constructed double chamber electrolysis cell)

### Coupling of the MEC and MFC

The anode of the MFC was connected to the cathode of the MEC and the cathode of the MFC was connected to the anode of the MEC as shown in Figure 4.

**Figure 4.**
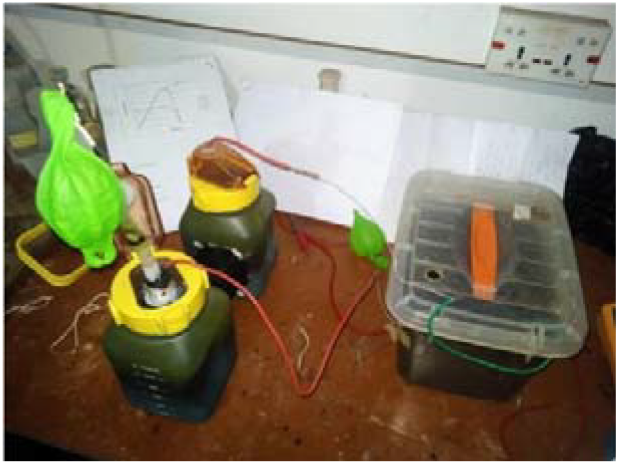
(An image of the MFC-MEC coupled system)

## Results and Discussion

### Results

After 13 days of the experiment, the MFC produced an average voltage of 118.9 ± 0.001 *mV* while the MEC produced an average voltage of 56.8 ± 0.003 *mV*. The coupled MEC-MFC system produced an average voltage of 87. ± 0.002 *mV* as well as an average current of 0.22 ± 0.006 *µA*. The average volume of hydrogen gas produced was about 6.9 ± 0.012 *mL/day*. The MFC produced a maximum of 310.7 ±*0.001mV* and the MEC also produced a maximum of 110.4 ± *0.003mV*, while the coupled system also produced a maximum of 155.9 mV. Also, the maximum hydrogen yield, which was approximately 65.5 ± 0.012 *mL/day* was on the 9th day of the experiment. In all of the experiment, a total amount of about 96.8 mL of hydrogen gas was produced and was tested using the pop test.

**Table 1.**
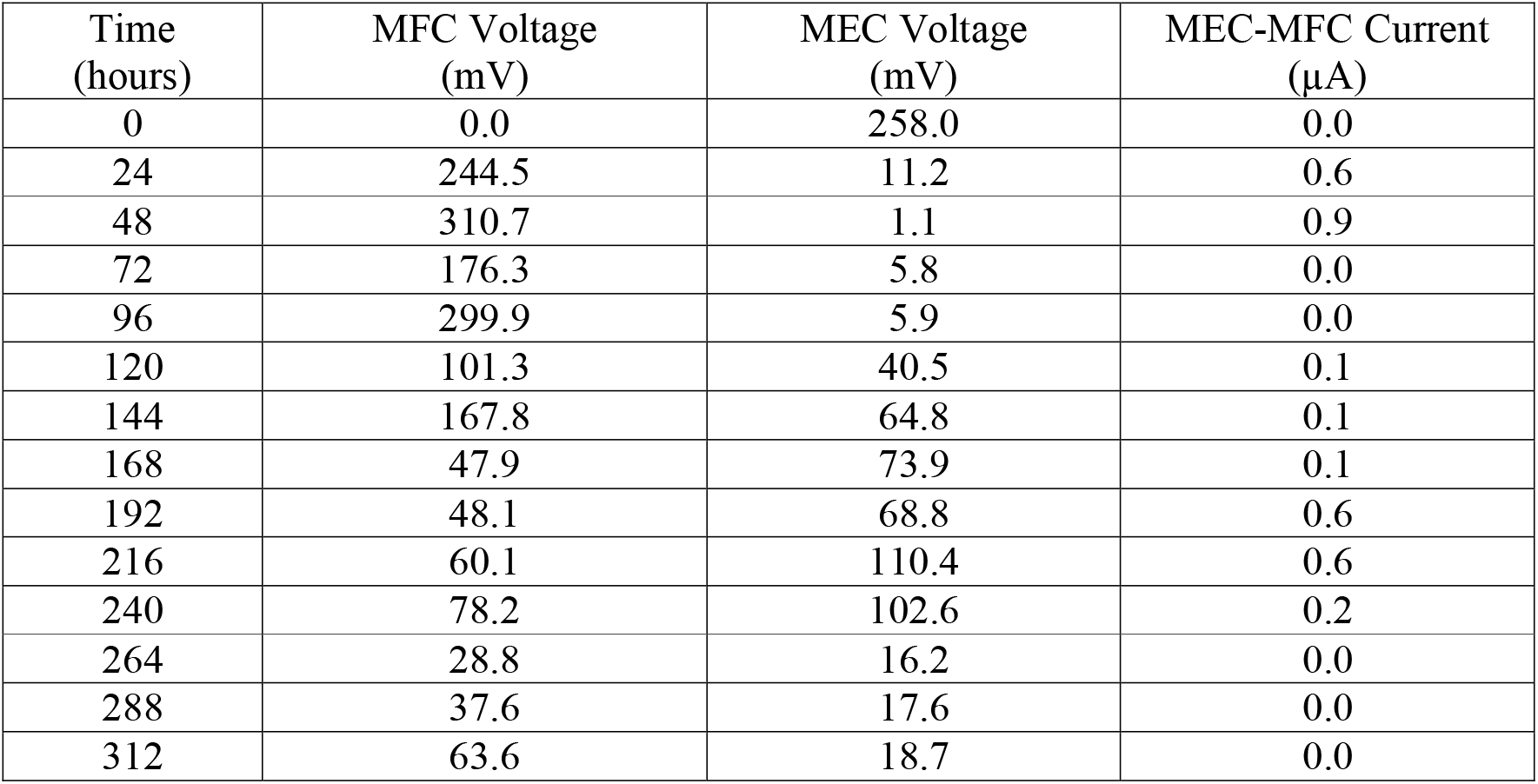
(Table of values for MFC and MEC voltages, current through the system and time)

| Time (hours) | MFC Voltage (mV) | MEC Voltage (mV) | MEC-MFC Current ($\mu\text{A}$) |
| --- | --- | --- | --- |
| 0 | 0.0 | 258.0 | 0.0 |
| 24 | 244.5 | 11.2 | 0.6 |
| 48 | 310.7 | 1.1 | 0.9 |
| 72 | 176.3 | 5.8 | 0.0 |
| 96 | 299.9 | 5.9 | 0.0 |
| 120 | 101.3 | 40.5 | 0.1 |
| 144 | 167.8 | 64.8 | 0.1 |
| 168 | 47.9 | 73.9 | 0.1 |
| 192 | 48.1 | 68.8 | 0.6 |
| 216 | 60.1 | 110.4 | 0.6 |
| 240 | 78.2 | 102.6 | 0.2 |
| 264 | 28.8 | 16.2 | 0.0 |
| 288 | 37.6 | 17.6 | 0.0 |
| 312 | 63.6 | 18.7 | 0.0 |

**Table 2.** (Table of values for time, daily radii of balloon, daily and cumulative volume of H_2_)

| Time (hours) | Daily radius of balloon (cm) | Daily Volume of H<sub>2</sub> (mL) | Cumulative Volume of H<sub>2</sub> (mL) |
| --- | --- | --- | --- |
| 0 | 0.0 | 0.0 | 0.0 |
| 24 | 0.0 | 0.0 | 0.0 |
| 48 | 0.0 | 0.0 | 0.0 |
| 72 | 0.0 | 0.0 | 0.0 |
| 96 | 0.0 | 0.0 | 0.0 |
| 120 | 1.3 | 9.2 | 9.2 |
| 144 | 1.7 | 20.6 | 29.8 |
| 168 | 0.5 | 0.5 | 30.3 |
| 192 | 0.3 | 0.1 | 30.4 |
| 216 | 2.5 | 65.5 | 95.9 |
| 240 | 0.6 | 0.9 | 96.8 |
| 264 | 0.0 | 0.0 | 96.8 |
| 288 | 0.0 | 0.0 | 96.8 |
| 312 | 0.0 | 0.0 | 96.8 |

### Discussion

Equation 1 was used to calculate the approximate volume of the hydrogen gas produced based on the radius (r) of the balloon recorded on each day of the experiment.

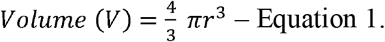

The coupling of the two systems allowed for electron transfer from the anode chamber of the MEC to the cathode of the MFC. Then, these electrons, together with those produced by the MFC, were transferred to the cathode chamber of the MEC for recombination.

It takes a while for exoelectrogens to form a film on the anode and also consume the nutrients from the organic matter to be able to give off electrons. This accounts for the reason why hydrogen gas was produced after about 120 hours as indicated by Figure 6. However, for the first few days where no quantified hydrogen gas was produced, it was seen that a lot of small bubbles had formed inside the anolyte solution in the cathode chamber of the MEC, symbolizing the formation of hydrogen gas.

Observing Figure 5, it is realized that there are two peaks with the highest being the first one, which implied that the current flowed highest on those two days of the experiment. It also projected that two peaks would be formed in Figure 6 since higher currents would lead to higher hydrogen production rates. Unfortunately, it is not seen as such. The second peak of Figure 5 aligns with that of Figure 6, which accounts for the highest rate of hydrogen production but not for Figure 6. For the first peak in Figure 5, it was observed on those days that several bubbles had formed in the cathode chamber. Those bubbles were made of hydrogen gas which had not risen into the balloon to be quantified.

**Figure 5.**
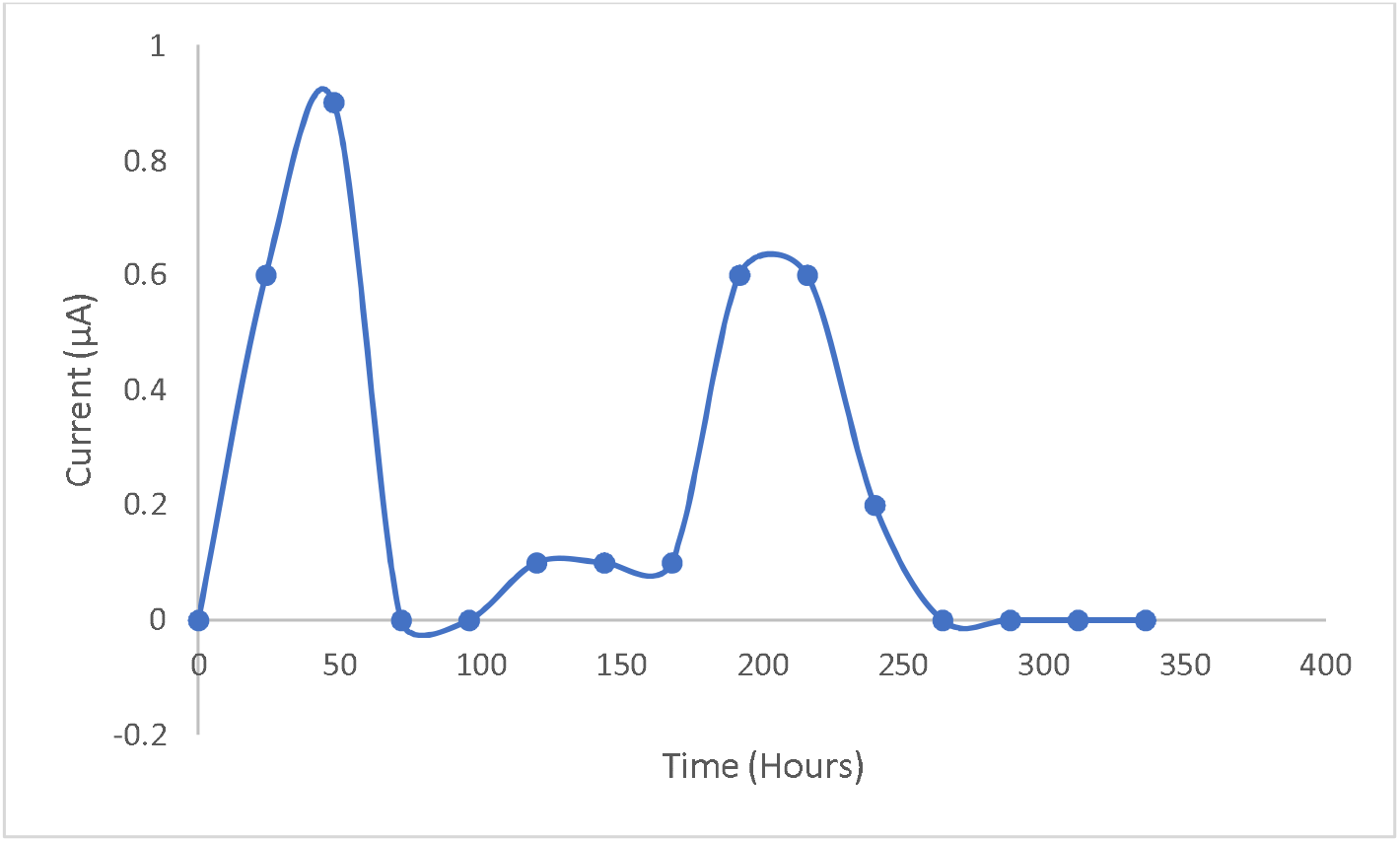
(A graph representing the rate of current production in the MEC-MFC coupled system)

**Figure 6.**
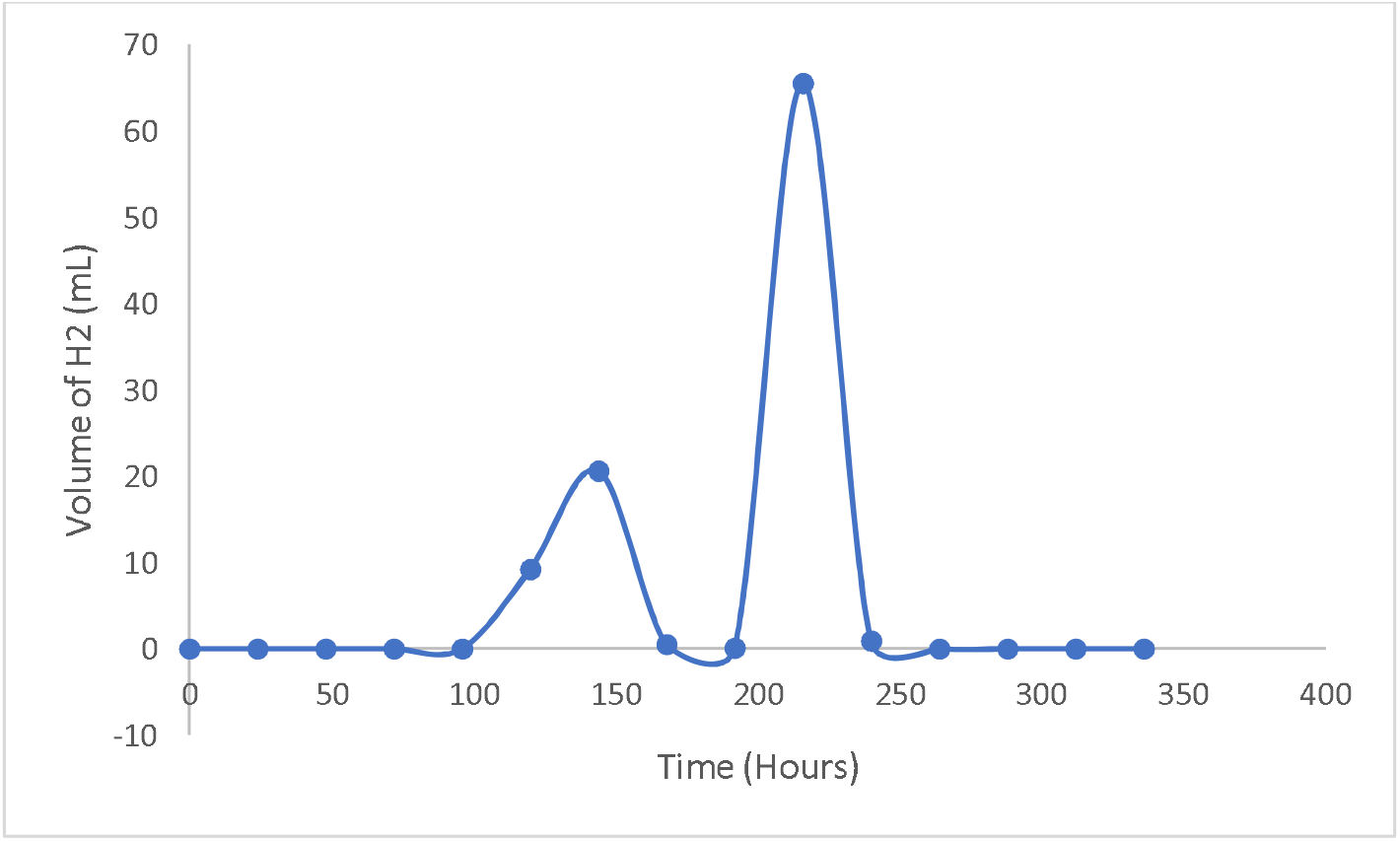
(A graph indicating the rate of hydrogen production in the MEC-MFC coupled system)

Also from Figure 5, it is seen that the current of the system dropped from the first peak to zero and recovered gradually, which proves that the microorganisms were already producing electrons when they were taken from their original habitation to be used for our work. The drop and gradual rise in current also meant that the microorganisms were adapting to the new system to be able to colonize the electrode, feed on the new source of nutrient, and release hydrogen ions and electrons which we needed in our work. This completely explains why the current of the coupled system kept fluctuating for some time. It kept fluctuating as the days went by until the 11th day, where it dropped to zero and maintained it for the remaining days of the experiment. This zero current recorded from day 11 to the end of the experiment implied that electrons were no longer being transported into the cathode chamber of the MEC for recombination to occur.

The complete drop in the hydrogen gas production from the 10th day to the end of the experiment also implies that the microorganisms could not get access to the nutrients in the organic matter to be able to feed and give off electrons as well as hydrogen ions.

As expected, the recorded average voltage of the MEC, 56.8 ± 0.003 mV, was very low. MECs are expected to produce lower voltages as compared with MFCs due to the purposes they each serve. In a case where the voltage across the MEC is greater than that of the MFC in a coupled system, a reverse reaction is expected to take place. This therefore would cause the MEC to act as an MFC, failing its purpose as an electrolysis cell.

## Conclusion and Recommendation

### Conclusion

The results of this work offer essential and new information on how feasible this technology is, for producing green hydrogen. A coupled MEC-MFC configuration with an average voltage of 87.9 ± 0.002 *mV* and an average current of 0.22 ± 0.006 *µA* was produced by the integration of these two systems. According to these data taken, the MFC was more effective at producing electricity than the MEC, which is in line with previous research that emphasizes the higher power densities that are generally linked to MFCs because of their capacity to directly utilize organic substrates through microbial metabolism. The maximum voltage outputs recorded were notably high. In terms of hydrogen production, the system produced 6.9 ± 0.012 *mL* of hydrogen gas on average every day, for a total yield of 96.8 mL for the course of the experiment. The highest hydrogen production ever measured was on day nine, at 65.5 ± 0.012 mL/day. This suggests the presence of optimal operating conditions or substrate availability that affects hydrogen generation rates at specific times. These results highlight the potential of MEC-MFC systems to support clean fuel alternatives by generating green hydrogen via biological processes that use biodegradable waste materials as substrates for microbiological activity. In addition, this research emphasizes important features like scalability and operational stability for potential uses in the renewable energy space, where hydrogen can be used as a flexible energy source or feedstock for a range of domestic and industrial operations.

### Recommendations

1. It is suggested that the system’s design and operational characteristics could be modified to improve the relatively low voltages and currents.
2. Future research on this work should consider adding more of the source of nutrients to allow the system to run for a longer period of time.
3. More investigations should be done to improve the efficiency and practicality in real-world situations, even though more optimization is required to optimize both voltage output and hydrogen production rates in coupled MEC-MFC systems.

